# *In vivo* optogenetic stimulation of the primate retina activates the visual cortex after long-term transduction

**DOI:** 10.1101/2021.02.09.427243

**Authors:** Antoine Chaffiol, Matthieu Provansal, Corentin Joffrois, Kévin Blaize, Guillaume Labernede, Ruben Goulet, Emma Burban, Elena Brazhnikova, Jens Duebel, Pierre Pouget, José Alain Sahel, Serge Picaud, Gregory Gauvain, Fabrice Arcizet

## Abstract

Over the last 15 years, optogenetics has changed fundamental research in neuroscience, and is now reaching toward therapeutic applications. Vision restoration strategies using optogenetics are now at the forefront of these new clinical opportunities. But applications to human patients suffering from retinal diseases leading to blindness rise important concerns on the long-term functional expression of optogenes and the efficient signal transmission to higher visual centers. Here we demonstrate in non-human primates, continued expression and functionality at the retina level ∼20 months after delivery of our construct. We also performed *in-vivo* recordings of visually evoked potentials in the primary visual cortex of anaesthetized animals. Using synaptic blockers, we isolated the *in-vivo* cortical activation resulting from the direct optogenetic stimulation of primate retina.

In conclusion, our work indicates long-term transgene expression and transmission of the signal generated in the macaque retina to the visual cortex, two important features for future clinical applications.

## Introduction

Repairing sensory impairments has always been an overarching goal in medicine. In the particular case of vision loss, considerable progress has been achieved in recent years through the development of various therapeutic strategies, such as retinal prostheses (1-4), stem cell transplantation (5-8) and optogenetic therapies (9-19). All these approaches aspire to restore retinal visual information. Microbial opsin-based optogenetics is one of the most promising of these approaches. It involves the expression of light-sensitive ion channels in preserved inner retinal neurons, restoring the intrinsic light sensitivity of the pathological retina in several types of ocular disease.

In inherited forms of retinal degeneration, such as retinitis pigmentosa (RP), the retinal degeneration is progressive, beginning with the photoreceptors and inevitably leading to blindness (20). The choice of target cell type in the retinal circuit should take into account the potential for translation into clinical applications and uses in patients. The accessibility of the targeted cell population, and the maintenance of its structure and integrity after the onset of retinal degeneration are key features. Since the first use of optogenetics to restore vision in blind mice through the expression of channelrhodopsin-2 (Chr2) in RGCs (9), many other studies have been conducted, targeting different cell types in the retina: photoreceptors (10, 21), bipolar cells (15, 16, 18) or retinal ganglion cells (RGCs) (9, 11, 13, 14, 17). Importantly, in diseases such as age-related macular degeneration (AMD) and RP, the retinal ganglion cells (RGCs) remain well-preserved during the process of retinal degeneration, even at late stages of the disease, after the death of the photoreceptors (22). Various models, including rodents, non-human primates, post-mortem human retina and human induced pluripotent stem cells (hIPS), have been used for investigations of optogenetic approaches, with promising results (7, 9, 13, 21).

Primate models seem to be the best of the animal models for testing optogenetic therapeutic approaches, because they share essential anatomical features and a similar organization of visual pathways with humans (23, 24). However, few studies to date have used this animal model for investigations of the potential of optogenetic therapy for the retina. For example, several opsins targeting RGCs have been tested in *ex vivo* preparations, including the microbial opsin channelrhodopsin 2 (14) in marmosets and CatCh (11), ReaChr (19) and ChR-tdT (13) in macaques. All these opsins were found to be functional in RGCs. Furthermore, the optogenetic responses of RGCs *in vivo* have been recorded with calcium imaging after photoablation of the photoreceptors in the macaque foveal region (17). However, none of these experiments was able to demonstrate the transfer of optogenetic activation from RGCs to higher visual centers, such as the primary visual cortex. Such experiments on retinal optogenetic approaches have been formed only in rodents, and have shown that the optogenetic activation of transduced retina induces specific visual evoked responses (VEPs) in the visual cortex (10, 11, 15, 25, 26). Moreover, specific cortical responses were recorded following the activation of RGCs by photovoltaic subretinal implants in rats (27). A recent study on primates fitted with subretinal implants showed that these prostheses induced some behavioral responses (2). However, to our knowledge, no study to date has demonstrated the transmission of information to higher visual areas following activation of the transduced retina in primates.

Using this approach to optogenetic therapy, we targeted the retinal ganglion cells (RGCs) in primate retinas through the *in vivo* expression of an ectopic light-sensitive ion channel, ChrimsonR, coupled to the fluorescent reporter tdTomato (13). The possible application of this strategy to blind patients suffering from retinal dystrophies raises important concerns about long-term functional expression to ensure efficient signal transmission to higher brain centers (i.e. the visual cortex) (28). We previously showed that the transduced retina displays a high degree of spatiotemporal resolution *ex vivo*, compatible with the perception of highly dynamic visual scenes at light levels suitable for use in humans (13). Here, we demonstrate, in non-human primates, sustained functional efficacy ∼20 months after the delivery of an AAV2.7m8-ChrimsonR-tdTomato vector similar to that currently undergoing clinical evaluation. Our results reveal a persistence of expression in the perifovea, mediating information transfer to higher brain centers. Indeed, we recorded visual evoked potentials in the primary visual cortex of anesthetized primates in response to optogenetic retinal activation. We used an intravitreal injection of synaptic blockers to isolate the cortical component resulting from the *in vivo* optogenetic stimulation of primate RGCs. Our findings demonstrate the long-term functional efficacy of optogenetic therapy to restore information transfer from the retina to the brain *in vivo*.

## Results

The experiments were conducted on three monkeys (*Macaca fascicularis*); each of them receiving, in one eye, a single intra-vitreal injection of AAV2.7m8-ChrR-tdT at a dose of 5 × 10^11^ vg/eye, the other eye being kept as a control. More than 20 months later, we performed *in-vivo* cortical experiments to record visually evoked potentials to the retinal optogenetic stimulation. Subsequently, ∼24h hours after the euthanasia, we measured directly optogenetic responses on *ex-vivo* retinal foveal explants. For every eye treated with ChrR-tdT, half of the retina was used for single-cell RGC recordings and 2-photon imaging, whereas the other half was used for Multi Electrode Array (MEA) RGCs population recordings and later histology.

The long-term functional expression of ChrimsonR-tdTomato (ChR-tdT) was assessed in the primate retina by *ex vivo* retinal recordings in the presence of glutamate receptor antagonists (see Methods), which were added to the bath solution to suppress any natural light response. Live epifluorescence images revealed a high density of transfected cells localized in the perifoveolar region, forming a torus shape (**Fig. 1A**), mostly in the retinal ganglion cell layer (**Fig. 1B, Sup. Fig. 1**). On some of these transfected RGCs, we observed small dendritic arbors suggesting that foveal midget RGCs responsible for high acuity vision (29) were expressing ChR-tdT. Light sensitivity and temporal dynamics were measured, at the single-cell level, with the two-photon guided patch-clamp technique, on ChR-tdT-positive cells (**Fig. 1C**). We found that the mean normalized photocurrent response increased significantly with increasing light intensity (**Fig. 1D**, top), reaching a peak (mean of 143.6 ±47.4 pA, *n*=12) at a light intensity of 3 × 10^17^ photons.cm^-2^.s^-1^. In the loose-patch recording configuration, firing rate displayed a similar dependence on light level (99.03 ±7.59 Hz, *n*=32 cells), also peaking at a light intensity of 3 × 10^17^ photons.cm^-2^.s^-1^ (**Fig. 1E, right**). We then showed that the highest frequency responses were obtained for light stimuli at 575 nm (**Fig. 1F**), corresponding to the excitation peak of ChrimsonR (590 nm) (30). We obtained reliable firing bursts with fast dynamics during the measurement of RGC responses to stimuli of increasing durations (20 ms to 4 s, *n*=9, **Fig. 1G**) or various flicker frequencies (10 repeats in full duty cycle) up to 28 Hz (*n*=9, **Fig. 1H**). These results demonstrate the ability of these engineered cells to follow and resolve short, long and fast light stimulations accurately up to frequencies very similar to the video-rate frequencies (∼25 Hz) required for fluid movement perception and compatible with the limits for flicker perception reported for humans (31).

**Figure 1.**
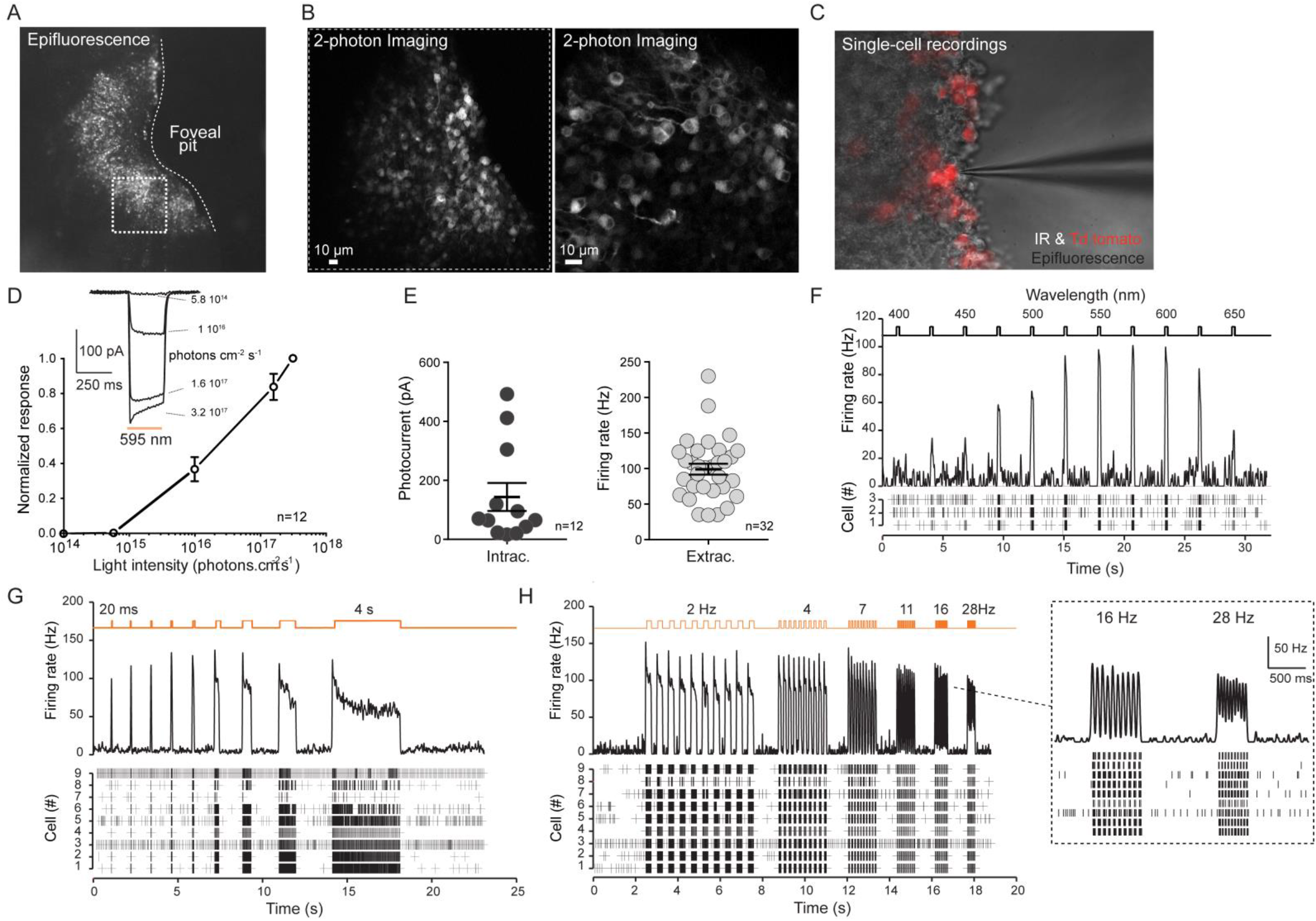
Long-lasting expression of the optogene in the macaque perifoveal ring and two-photon guided single-cell recordings. (A) Epifluorescence image of an *ex vivo* primate hemifovea expressing tdTomato-ChrimsonR (perifoveal ring) 20 months after the injection of AAV2.7m8 -ChR-tdT at a dose of 5 × 10^11^ vg/eye (B) Two-photon images of tdTomato-positive cells in the retinal ganglion cell layer at two different magnifications, taken in the dotted square shown in (A). (C) Combined infrared and epifluorescence image of tdTomato-expressing fluorescent cells during single-cell recordings with a glass electrode (patch-clamp or extracellular recordings). (D) Photocurrent response as a function of the intensity of the light stimulus in transfected retinal ganglion cells 20 months post-injection, and comparison with earlier time points. (top) Example of photocurrents recorded in cells stimulated with light at 595 nm for 250 ms at intensities of 5.8 × 10^14^ to 3.2 × 10^17^ photons.cm^2^.s^-1^. (bottom) Normalized intensity curve for 12 patched cells. (E) Peak responses for photocurrent (left, patch-clamp data, *n* = 12 cells) and firing rate (right, extracellular recordings, *n* = 32 cells) recorded in RGCs for a light stimulus at a wavelength of 595 nm and an intensity of 3.2 × 10^17^ photons.cm^2^.s^-1^. (F) Spectral tuning. RGC response (raster plot and firing rate) as a function of the stimulus wavelength tested, for wavelengths between 400 and 650 nm (*n* = 3 cells). The peak response (asterisk) was observed at 575 nm. The black curve represents the mean cell response. (G) RGC response as a function of stimulus duration. Raster plot (bottom) for nine cells and firing rate (top), with stimuli of increasing duration, from 20 ms to 4 s. The black curve represents the mean cell response. (H) Temporal resolution. RGC activity as a function of stimulation frequency, represented as a raster plot (bottom) and firing rate (top), for 10 flicker repeats at frequencies from 2 Hz to 28 Hz (*n* = 9 cells). The black curve represents the mean cell response.

In parallel, we used MEA recordings to investigate the responsiveness of the cell population in the transfected area (**Fig. 2, Sup. Fig. 2**). A large proportion of the perifovea contained a high density of ChR-tdT-expressing cells, as indicated by counting TdTomato-positive cells on projections of confocal stacks (**Fig. 2A-B**). The MEA chip (**Fig. 2C**, top left) covered a large area of the hemifoveal retina flat mounts, making it possible to take recordings for a large proportion of the transfected RGCs (**Fig. 2C**, bottom left; **Sup. Fig. 2A**). **Figure 2C** (right) and **Sup. Fig. 2C** show the global recorded activity for RGCs, represented as the firing rate of 256 sites over a period of 4 s in response to a 2 s flash of light. We performed 10 recordings in total, at a wavelength of 595 nm and a light intensity of 7 × 10^16^ photons.cm^2^.s^-1^ (**Fig. 2C**, right). A large proportion of the recording sites were responsive, and the response observed was correlated to the degree of optogene expression, as determined by measuring tdTomato reporter fluorescence. Overall, in the two retinas for which we were able to obtain spontaneous activity, 45.5% of all active recording sites were also responsive to light stimulation (172 of 378 active sites, **Fig. 2D**). We performed MEA recordings as a function of light intensity (**Fig. 2E**). The results obtained were very similar to those obtained with the single-cell technique (**Fig. 1D-E**). Indeed, the threshold light intensity for a response was found to be 9 × 10^15^ photons.cm^2^.s^-1^. We then analyzed MEA responses as a function of stimulus duration (**Fig. 2F**), at an intensity of 7 × 10^16^ photons.cm^2^.s^-1^. In the two retinas tested, cells responded to stimuli with a duration of at least 5 ms, but not to stimuli of shorter duration.

**Figure 2.**
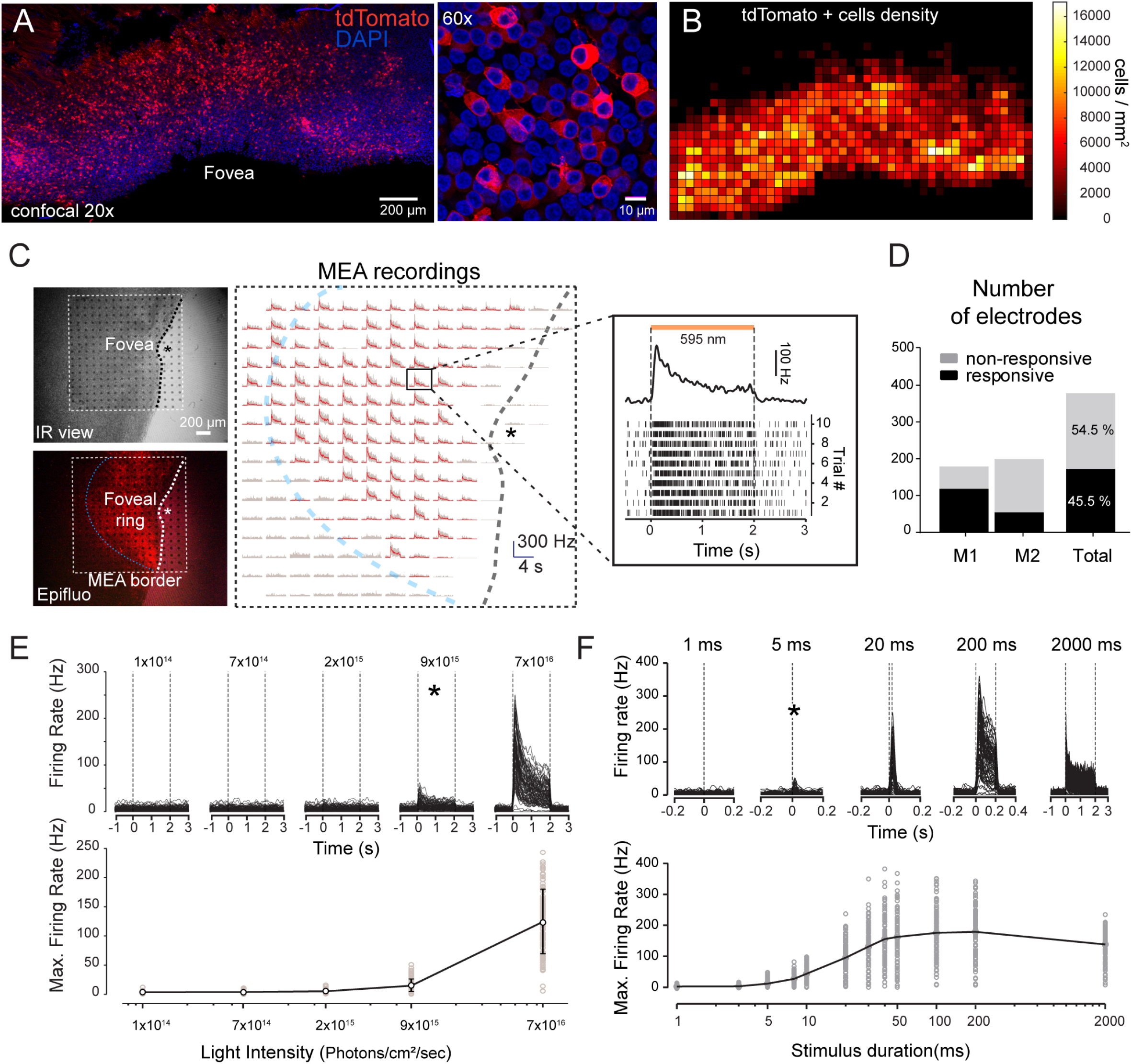
Expression and functionality: multi-electrode array recordings. (A) Projection of confocal stack stitches (20x objective) showing the perifoveal area of M1 retina 20 months after treatment with a vector dose of 5 × 10^11^vg/eye (left). ChR-tdT-expressing cells are shown in red, and the nuclei (in blue) are stained with DAPI. Scale bar: 200 µm. (Right) Higher magnification image (60x objective). Scale bar: 10 µm. (B) Density map of ChR-tdT-positive RGCs for the hemifovea shown in (A). (C) Hemifovea RGC population recordings with a MEA (monkey M1). *Top left*, infrared view of the hemifovea with a 256-site MEA placed on the side of the RGC layer. *Bottom left*, epifluorescence image of the same retina showing tdTomato-ChrimsonR fluorescence. *Right*, global representation of the recorded RGC activity (firing rate) of the 256 sites during a 4 s period, with the application of a 2 s flash of light (at a wavelength of 595 nm). *Right, inset*, Raster plot (bottom) and spike-density function (top) of a representative recording site with 10 repeats. The dotted squares represent the MEA borders, the blue curves represent the approximate limits of Chrimson-tdTomato expression (half-donut shape) and the asterisks represent the center of the fovea. (D) Proportion of responsive versus non-responsive MEA electrodes for the retinae of the two primates (NHPs M1 and M2). (E) Firing rate (top) of 118 responsive electrodes (monkey M1 shown as an example) as a function of light stimulus intensity (black curves) and mean response intensity curve for these neurons (bottom). The asterisk represents the threshold light intensity for the elicitation of light responses. Light intensities of between 1 × 10^14^ and 7 × 10^16^ photons.cm^2^.s^-1^were tested. (F) Firing rate (top) of RGCs shown in (F) as a function of stimulus duration (black curves), and the mean population response curve (bottom). The asterisk represents the threshold light duration required to elicit light responses. Stimulus duration ranged from 1 ms to 2000 ms.

In living animals, we then investigated whether the optogenetic stimulation of these transfected RGCs could activate the visual pathway *in vivo*. To this end, we recorded visual evoked potentials (VEPs) in response to different visual stimuli presented to the anesthetized animals during short-term experiments (**Fig. 3A**). In this way, we were able to compare VEP responses before and after the intravitreal injection of synaptic blockers (PDA and L-AP4) into both eyes (**Fig. 3B**) to block natural retinal responses to light (32). With the synaptic block, light stimulation directly activates the engineered cells rather than the normal retinal pathway. In monkey M1, following stimulation of the ChR-tdT-expressing eye with an orange LED (595 nm) before the injection of the blocker, we recorded typical VEP responses, defined by the presence of four different phases: two negative phases with latencies of 28 and 76 ms (blue triangles) and two positive phases (latencies of 51 and 107, red triangles). Similar VEPs were obtained for stimulation of the control eye, confirming that the sequence of orange flashes activated the visual pathway naturally. After the intravitreal injection of synaptic blockers and stimulation of the eye with the same orange LED, we noted major changes in the shape of the recorded VEP. The VEP responses of the ChR-tdT-expressing eye (**Fig. 3B** orange traces) were characterized by an early positive phase (21 ms, orange arrow) and a late sustained negative phase (101 ms, blue arrow). Interestingly, stimulation of the control eye resulted in a stable recording, with no early positive or subsequent negative phase. Similar results were obtained with a second animal (monkey M2, **Fig. 3B bottom**), with an early peak response at 24 ms. The latency of the first positive peak was 21-24 ms for both animals, which is much shorter than the latencies classically reported for natural VEP components (positive components occur at 50-81 ms) and observed here. These data are consistent with the notion that these VEP responses in the visual cortex were elicited by the direct activation of ChR-tdT-expressing RGCs.

**Figure 3:**
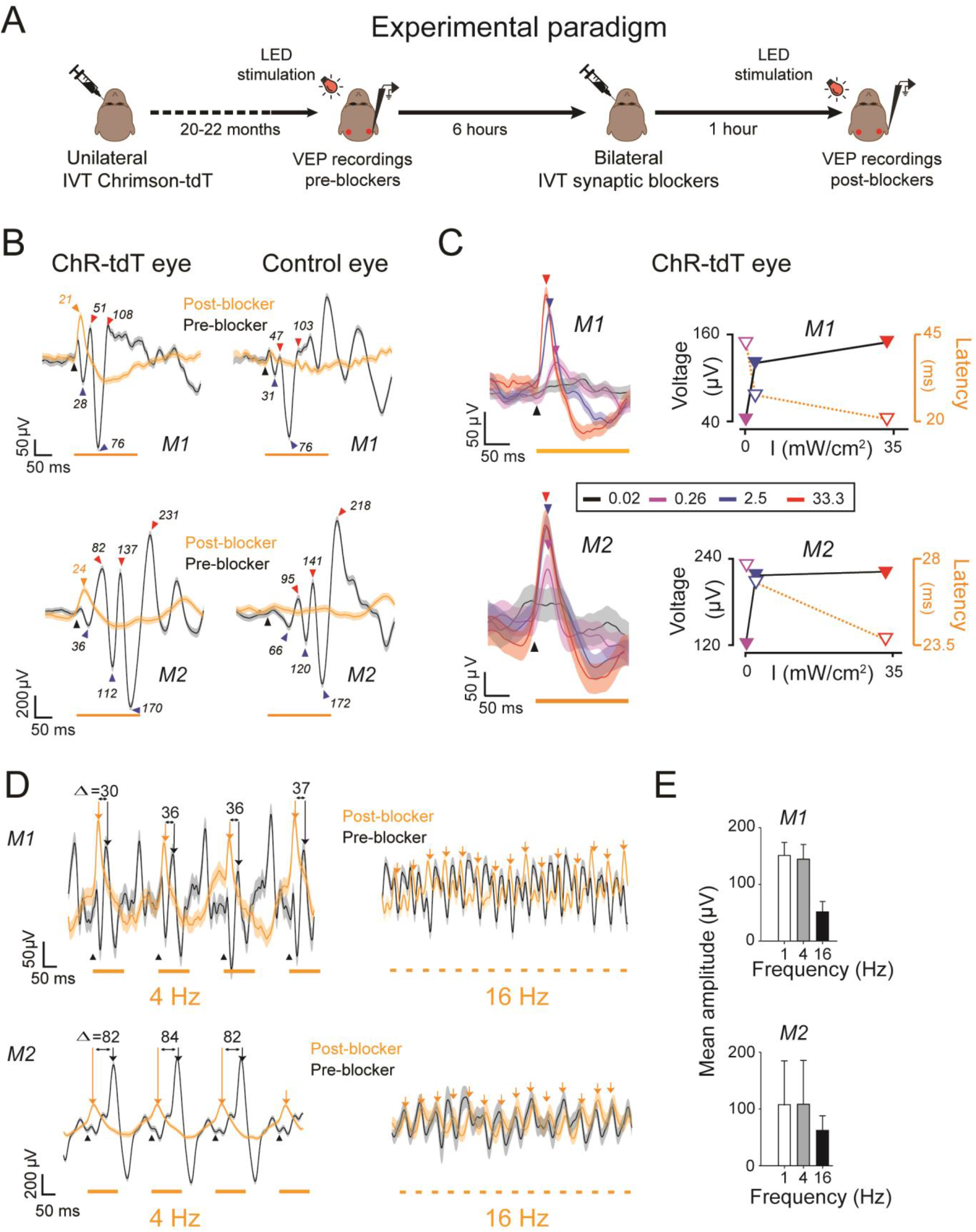
Cortical responses after retinal ChrimsonR optogenetic stimulation. (A) Experimental set-up. Both animals received injections of AAV-ChrimsonR-Tdtomato into one eye. We recorded VEP responses 20-24 months after the injection, following various protocols of LED flash stimulation of the anesthetized animal. We then injected a synaptic blocker to block photoreceptor transmission. Finally, we recorded VEP responses for the same LED flash stimulation protocols. (B) VEP responses to the orange LED before (black curve) and after (orange curve) the injection of synaptic blocker. The left column shows VEP responses following stimulation of the ChR-Tdt eye, and the right column shows the response to stimulation of the control eye. VEP responses are shown in the top row for monkey M1 and the bottom row for monkey M2. Black inverted triangles indicate the start of LED stimulation, orange triangles indicate the peak latencies of VEP responses after treatment with a synapse blocker. Red triangles indicate positive VEP peaks before injection, and blue triangles indicate the negative phases. Numbers indicate peak latencies in ms. (C) VEP responses to stimulation of the ChR-Tdt eye with different light intensities and frequencies. We tested four different intensities at a frequency of 1 Hz (33.3, 2.5, 0.26 and 0.02 mW/cm^2^), as indicated by the color code. (D) We also tested two different light frequencies (4 and 16 Hz). We report the differences between peak latencies in ms (e.g. Δ30). Orange arrows indicate positive phases of VEP responses. (E). Mean amplitudes of positive phases of VEP responses for different light frequencies (1, 4 and 8 Hz) for monkey 1 (*top, M1*) and monkey 2 (*bottom, M2*). Errors bars indicate SEM.

We then investigated the effects of several parameters, such as the light intensity and frequency of the orange LED, on VEP responses (**Fig. 3C)**. The amplitude of positive peak responses increased proportionally to light intensity (from 43 to 152 μV for monkey 1 and from 121 to 221 μV for monkey 2), and the peak latencies of these responses decreased with increasing light intensity (see inset); from 82 to 24 ms for monkey M1 and 28 to 24 ms for monkey M2). No significant VEP response was recorded for the control eye (**Sup. Figure 3**), even at maximal light intensity, confirming that the glutamate receptor antagonists used effectively abolished the natural light response. We then recorded VEP responses in response to an orange LED flashed at a frequency of 4 Hz (125 ms) and 16 Hz (32 ms, **Fig. 3C)** to confirm that activation occurred earlier after the optogenetic stimulation of RGCs. VEP peaks occurred more rapidly after blocker administration (orange traces) than in the absence of blocker (black traces), with similar time periods observed for stimulation at 4 Hz and at 1 Hz (30-37 ms for monkey M1; Fig 3B). For monkey M2, the difference in time to VEP peak was larger at 4 Hz (82-84 ms) than at 1 Hz (∼58 ms). After stimulation at 16 Hz, VEP traces followed the train of pulses both before and after the injection of synaptic blocker. Although mean amplitudes of VEP responses were similar for stimulations at 1 Hz and 4 Hz, they decreased drastically after stimulation at 16 Hz for both animals (Fig. 3E). No such activation was observed following stimulation of the control eye, in either of the animals studied (Sup. Fig. 3). These results demonstrate that optogenetic activation of the retina can trigger a transfer of information to higher visual centers, providing additional support for the potential of ChR-tdT for future therapeutic applications.

## Discussion

We previously showed that a single intravitreal injection of AAV2.7m8.CAG.ChrimsonR-tdT can efficiently target the foveal region of the retina, which is responsible for high-acuity vision (13). Using *ex vivo* live imaging and histology, we show here that expression persists in cells in the perifovea region more than 20 months after the injection, and we demonstrate, with electrophysiological recordings, that these transfected RGCs remain functional, displaying rapid, robust responses. Previous studies based on *ex vivo* retinal recordings in NHP models have demonstrated efficient optogene expression for up to six months (11, 13, 14), and a study based on retinal imaging *in vivo* extended this functional window up to 14 months (17). This maintenance of activity so long after the injection is consistent with the notion that gene therapy can lead to long-term gene expression. Furthermore, it provides additional evidence that the microbial opsin ChR-tdT does not induce an immune response that might eventually destroy the engineered RGCs. Indeed, the RGC responses were in a range similar to that recorded in our initial experiments (13), and transfected cell density was also at similar levels. However, given the small number of replicates in this study and the considerable inter-subject variability previously observed after two or six months, it is difficult to make quantitative comparisons with earlier expression time points.

We show here, for the first time, through *in vivo* VEP recordings, that the selective stimulation of transfected RGCs induces specific cortical responses. Based on our retinal observations, we can interpret these VEP recordings as reflecting the activation of cortical neurons due to the direct functional optogenetic activation of RGCs, leading to the synaptic transfer of information to cortical neurons. All previous studies on optogenetic RGC activation in the primate retina were performed on RGCs either *ex vitro* (11, 13, 14) or *in vivo* (17). The recorded rates of RGC firing activity and the reported increases in calcium indicator fluorescence were highly suggestive of potential information transfer to the higher visual centers, but no experimental demonstration for the existence of this communication was provided. Here, following a blockade of glutamatergic synaptic transmission in the retina, the direct optogenetic activation of RGCs elicited specific VEP responses with an earlier peak response than normal VEP responses. Such early cortical responses were recorded in blind rodents during optogenetic activation of either the dormant cones (10), the bipolar cells (15, 16) or the RGCs(11, 26). The short VEP response latencies recorded are probably the signature of direct optogenetic stimulation within the inner retina, occurring more rapidly than natural responses, due to the slowness of the phototransduction cascade and of synaptic information transfer between the different retinal layers. We showed that cortical VEP responses increased with light intensity. This result is highly consistent with the RGC spike recording on the isolated retina, with a clear relationship between RGC activity and light intensity. Given the high light levels used in our VEP experiments, the observed increases in VEP peak amplitudes with increasing light intensity are consistent with an optogenetic origin, because any residual natural light responses would be fully saturated at such light intensities. Finally, these VEP recordings show that optogenetic responses can follow frequencies of at least 16 Hz, as expected from the high temporal resolution achieved with RGCs in *ex vivo* single-cell recordings (Fig. 1H). All these VEP recordings validate the therapeutic potential of ChR-tdT for restoring vision in blind patients.

In conclusion, this study extends existing data for vision restoration strategies tested in NHPs, and raises hopes of long-term functionality for optogenetic approaches in blind patients, who may benefit from this therapy. It also opens up new avenues of research into the neural integration and computations occurring at the cortical level in NHPs, with a view to restoring the sensitivity of sensory organs through optogenetics.

## Materials and Methods

### Animals

Data were collected for three captive-born macaques (*Macaca fascicularis*; 2 males, monkey M1, monkey M2, weighing 3.2, and 3.9 kg, respectively; one female, monkey M0, 4.1 kg). Monkeys were housed in pairs and handled in strict accordance with the recommendations of the Weatherall Report on good animal practice. Monkey housing conditions, surgical procedures and experimental protocols were performed in strict accordance with the National Institutes of Health Guidelines (1996), and after validation of the European Council Directive (2010/63/EU), and the study was approved by the French government and institutional and regional committees for animal care (Committee C. Darwin, registration #9013). Our routine laboratory procedures included an environmental enrichment program, in which the monkeys were allowed visual, auditory and olfactory contact with other animals and, when appropriate, could touch and groom each other.

### AAV production

ChrimsonR-tdTomato was inserted into an AAV backbone plasmid. The construct included a WPRE and bovine growth hormone polyA sequences. Recombinant AAVs were produced by the plasmid cotransfection method (32), and the resulting lysates were purified by iodixanol gradient ultracentrifugation, as previously described. Briefly, the 40% iodixanol fraction was concentrated and subjected to buffer exchange with Amicon Ultra-15 Centrifugal Filter Units. Vector stocks were then titered for DNase-resistant vector genomes by real-time PCR relative to a standard (33).

### Gene delivery

Primates were anesthetized with 10:1 mg/kg mixture of ketamine and xylazine. We injected 100 µL of viral vector suspension into the vitreous of one eye in each animal. Following the injection, an ophthalmic steroid and an antibiotic ointment were applied to the cornea. Experiments were conducted 21, 20 and 22 months after injection for M0, M1 and M2, respectively. None of the treated animals displayed any sign of photophobia or vision-related behavioral changes during housing.

### Recording of VEPs

We performed *in vivo* VEP recordings 20 to 22 months after AAV injection, during a terminal experiment in two animals (M1 and M2). Briefly, anesthesia was induced with ketamine (0.2 mg/kg, intramuscular: i.m.), and dexmedetomidine (0.015 mg/kg, i.m.) and maintained with alfaxan (0.1 mg/kg/, min, i.v.). The monkey was placed in a stereotaxic frame and heart rate, temperature, respiration, and peripheral oxygen saturation were monitored throughout the experiment. We placed four electrodes in subcutaneous positions: two at each temple and two at each occipital operculum (left and right), and we set the high- and low-path filters to 50 Hz and 0.05 Hz, respectively. Light stimuli were generated with two different LEDs: a blue LED (M470L3 Thorlab) and an orange LED (M595L3 from Thorlab 595 nm). VEP responses were recorded before and after the intravitreal injection of synaptic blockers — 2,3-piperidine dicarboxylic acid (PDA) and L-(+)-2-amino-4-phosphonobutyric acid (L-AP4) — used to block natural light responses (34). The eyes of the monkeys were subjected to flash stimuli at various intensities (from 0.02 to 33.3 mW/cm^2^) and frequencies (1, 4 and 16 Hz). The duration of the stimuli was 200, 125 and 32 ms for frequencies of 1, 4 and 16 Hz, respectively. We made 300 consecutive recordings of each VEP response and then averaged response waveforms for each VEP measurement. VEP responses were similar for the ipsilateral and contralateral electrodes. We therefore present VEP responses for the electrode contralateral to the injected eye only.

### Primate retina isolation and preservation

After the *in vivo* VEP recordings, the primates received a lethal dose of pentobarbital. Their eyeballs were removed, perforated with a sterile 20-gauge needle and placed in sealed bags with CO_2_ independent medium (Thermo Fisher scientific) for transport. The retinae were isolated, and the retinal pigment epithelium was removed and stored as retinal explants in an incubator for ∼24 hours before recording. Hemifoveal retina fragments were transferred to Neurobasal + B27 medium in polycarbonate Transwells (Corning) for conservation in the cell culture incubator. These hemifoveal regions were subsequently used for simultaneous single-cell and MEA recordings. In these conditions, natural photoreceptor responses were abolished and did not recover. We abolished all natural responses entirely, by applying pharmacological blockers (see below).

### Two-photon live imaging and single-cell electrophysiological recordings

A custom-built two-photon microscope equipped with a 25x water immersion objective (XLPLN25xWMP, NA: 1.05, Olympus) with a pulsed femtosecond laser (InSight™ DeepSee™ -Newport Corporation) was used for imaging ChR-tdT-positive retinal ganglion cells. AAV-treated macaque retinas were imaged in oxygenized (95% O_2_, 5% CO_2_) Ames medium (Sigma-Aldrich). For live two-photon imaging, whole-mount retinas (without the retinal pigment epithelium attached) were placed in the recording chamber of the microscope (ganglion cell layer side up), and images and *z*-stacks were acquired with the excitation laser at a wavelength of 1050 nm. Images were processed offline with ImageJ.

We used an Axon Multiclamp 700B amplifier for whole-cell patch-clamp and cell-attached recordings. Patch electrodes were made from borosilicate glass (BF100-50-10, Sutter Instruments) and pulled to 7-9 MΩ. Pipettes were filled with 115 mM potassium gluconate, 10 mM KCl, 1 mM MgCl_2_, 0.5 mM CaCl_2_, 1.5 mM EGTA, 10 mM HEPES, and 4 mM ATP-Na_2_ (pH 7.2). We clamped the cells at a potential of -60 mV, to isolate excitatory currents. Recordings were also performed in the loose-patch configuration with the pipettes filled with Ames medium, to record spiking activity. The retinae were dark-adapted for at least 30 minutes in the recording chamber before recordings. AMPA/kainate glutamate receptor antagonist, 6-cyano-7-nitroquinoxaline-2,3-dione (CNQX, 25 μM, Sigma-Aldrich), NMDA glutamate receptor antagonist, [3H]3-(2-carboxypiperazin-4-yl) propyl-1-phosphonic acid (CPP, 10 μM, Sigma-Aldrich) and a selective group III metabotropic glutamate receptor agonist, L-(+)-2-amino-4-phosphonobutyric acid (L-AP4, 50 μM, Tocris Bioscience, Bristol, UK) were diluted to the appropriate concentration from stock solutions and added to the Ames medium before recordings.

### MEA

Multielectrode array (MEA) recordings were obtained for retinal fragments (without the retinal pigment epithelium attached) placed on a cellulose membrane that had been incubated with polylysine (0.1%, Sigma) overnight. Once on the micromanipulator, the piece of retina was gently pressed against a MEA (MEA256 100/30 iR-ITO; Multi-Channel Systems, Reutlingen, Germany), with the retinal ganglion cells towards the electrodes. We measured tdTomato fluorescence to check that the retina was correctly positioned before making recordings under a Nikon Eclipse Ti inverted microscope (Nikon, Dusseldorf, Germany) with the MEA system mounted on the stage. The retina was continuously perfused with Ames medium (Sigma-Aldrich, St Louis, MO) bubbled with 95% O_2_ and 5% CO_2_ at 34 °C, at a rate of 1–2 ml/minute during experiments. AMPA/kainate glutamate receptor antagonist, 6-cyano-7-nitroquinoxaline-2,3-dione (CNQX, 25 μM, Sigma-Aldrich), NMDA glutamate receptor antagonist, [3H]3-(2-carboxypiperazin-4-yl) propyl-1-phosphonic acid (CPP, 10 μM, Sigma-Aldrich) and a selective group III metabotropic glutamate receptor agonist, L-(+)-2-amino-4-phosphonobutyric acid (L-AP4, 50 μM, Tocris Bioscience, Bristol, UK) were diluted to the appropriate concentration from stock solutions and added to the bath through the perfusion system for 10 minutes before recordings. Action potentials were identified on the filtered electrode signal (2^nd^-order high-pass Butterworth, cutoff frequency 200 Hz), with a threshold of 4 x the SD of the signal. Spike density function was calculated, averaged over repeat stimulations and used to determine the maximal firing rate over a time window corresponding to the duration of the stimulus plus 50 ms. In comparisons of responses between light intensities, we calculated, for each electrode, the added firing rate as the maximal firing rate minus the spontaneous firing rate for the electrode concerned, which was calculated as the mean firing rate in the 2 seconds before stimulation.

### Photostimulation for ex vivo experiments

For single-cell electrophysiological recordings, full-field photostimulation was performed with a Polychrome V monochromator (Olympus, Hamburg, Germany) set to 595 nm (± 10 nm), and output light intensities were calibrated and ranged from 5.8 × 10^14^ to 3.15 × 10^17^ photons.cm^2^.s^-1^. For spectral sensitivity experiments, stimulation wavelengths between 400 and 650 nm were tested, in 25 nm steps. For flicker stimulation, 10 repeats were used in full duty cycle, at frequencies ranging from 2 Hz to 28 Hz. For MEA recordings, full-field light stimuli were applied with another Polychrome V monochromator set to 595 nm (± 10nm), driven by a STG2008 stimulus generator (MCS). Output light intensities were calibrated, and ranged from 1.37 × 10^14^ to 6.78 × 10^16^ photons.cm^2^.s^-1^. For intensity curves, we used two-second flashes at five intensities (1.37 × 10^14^, 6.56 × 10^14^, 2.34 × 10^15^, 8.82 × 10^15^, 6.78 × 10^16^ photons.cm^2^.s^-1^), each repeated 10 times, with a five-second interval between stimuli. For the duration of stimulation assay, we used 12 different durations (ranging from 1 to 2000 ms), each repeated 10 times, with a five-second interval between stimuli. Calibrations were performed with a spectrophotometer (USB2000+, Ocean Optics, Dunedin, FL).

### Confocal imaging and quantification

After MEA experiments, the tissue was recovered and fixed by incubation with 4% PFA for 30 min at room temperature, rinsed with PBS and stored at 4 °C in sodium azide solution. Hemifoveas were then mounted in Vectashield containing DAPI (H-1000, Vector Laboratories) on slides and covered with a coverslip (18 x 18 mm, Biosigma), using a 100 µm spacer (Secure-seal space S24735, Thermo Fisher Scientific), which was sealed with nail polish. The retinas were imaged on an inverted confocal microscope (Fluoview 1200, Olympus), with a 20x objective (UPLSAPO 20XO, NA: 0.85, Olympus), voxel sizes of 0.265 to 0.388 µm/pixel in the *x* and *y* directions and 1.64 µm/pixel in the *z* direction. For each hemifovea, we recorded multiple stacks and reconstituted an automatic stitch (10% overlap). Using Td-tomato fluorescence, we performed manual 3D counts of the transfected cells in ImageJ (http://imagej.nih.gov/ij) with the cell counter plugin. The results were then processed with custom-developed matlab analysis software for the calculation of *local density*.

## Supporting information

Movie S1 (separate file). 2-Photon live imaging of a large volume of M1 hemifovea.

## Acknowledgments

We thank Morgane Weissenburger, Estelle Chavret-Reculon, Lucile Aubree, Corina Dussaud and Benedicte Daboval for their superb animal care. We thank Valérie Fradot for technical assistance with primate tissue preparation. This project was supported by BPIfrance (grant reference 2014-PRSP-15), Gensight Biologics, Foundation Fighting Blindness, *Fédération des Aveugles de France*, and by French state funds managed by the *Agence Nationale de la Recherche* within the *Investissements d’Avenir* program, RHU LIGHT4DEAF [ANR-15-RHU-0001], LABEX LIFESENSES [ANR-10-LABX-65], IHU FOReSIGHT [ANR-18-IAHU-0001], [ANR-11-IDEX-0004-02].

## Author contributions

Conceptualization, A.C., S.P., G.G. and F.A.; methodology & performance of experiments, A.C., M.P., C.J., K.B., G.L., R.G., E.Bu., E.Br., G.G. and F.A.; formal analysis, A.C., C.J., G.G. and F.A.; writing – original draft, A.C., F.A., G.G. and S.P.; writing – review & editing, A.C., F.A., G.G., P.P. and S.P.; supervision and funding, J-A.S., S.P., G.G. and F.A.

## Supplementary Data

Movie S1 (separate file). ***2-Photon live imaging of a large volume of M1 hemifovea***.

A 500 × 500 × 150 µm volume of M1 retina was imaged at an excitation wavelength of 1050 nm to visualize cells expressing the fluorescent reporter TdTomato. ImageJ and the 3Dviewer plugin were used to create a 360° video of the imaged area, revealing a high density of fluorescent cells in the RGC layer.

**Fig. S2.**
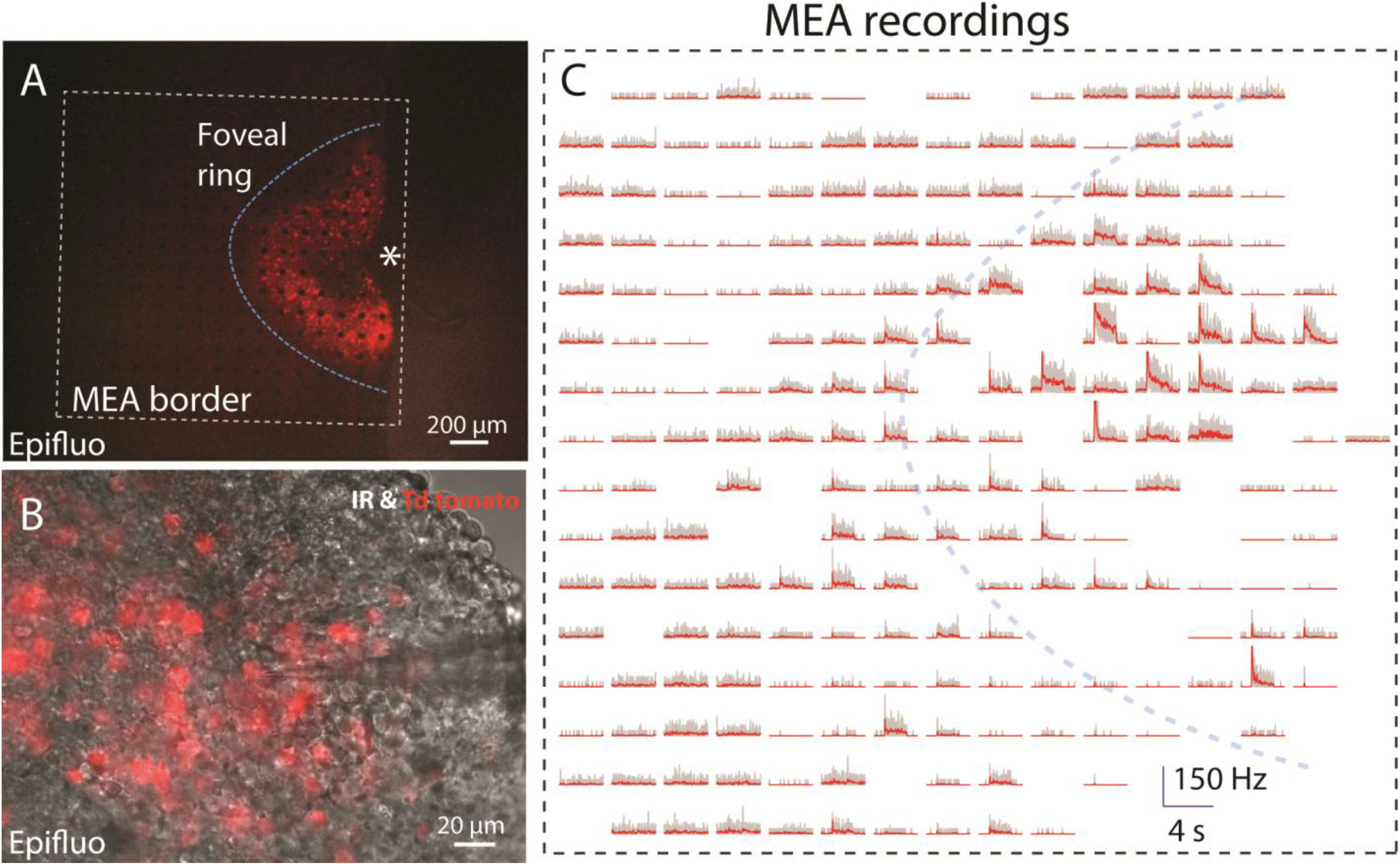
Expression and functionality: monkey M2. (A) Epifluorescence image of monkey M2 hemifovea showing tdTomato-ChrimsonR fluorescence, and with a 256-site MEA placed against the RGC layer. (B) Combined infrared and epifluorescence image of tdTomato fluorescent cells during single-cell recordings with a glass electrode (patch-clamp or extracellular recordings). Images were acquired from the second hemifovea. (C) Global representation of the recorded RGC activity (firing rate) for the 256 MEA recording sites shown in (A) over a period of 4 s, with the application of a 2 s flash of light (at a wavelength of 595 nm). The dotted squares represent the MEA borders, the blue curves represent the approximate limits of Chrimson-tdTomato expression (half-donut shape) and the asterisk represents the center of the fovea.

**Fig. S3.**
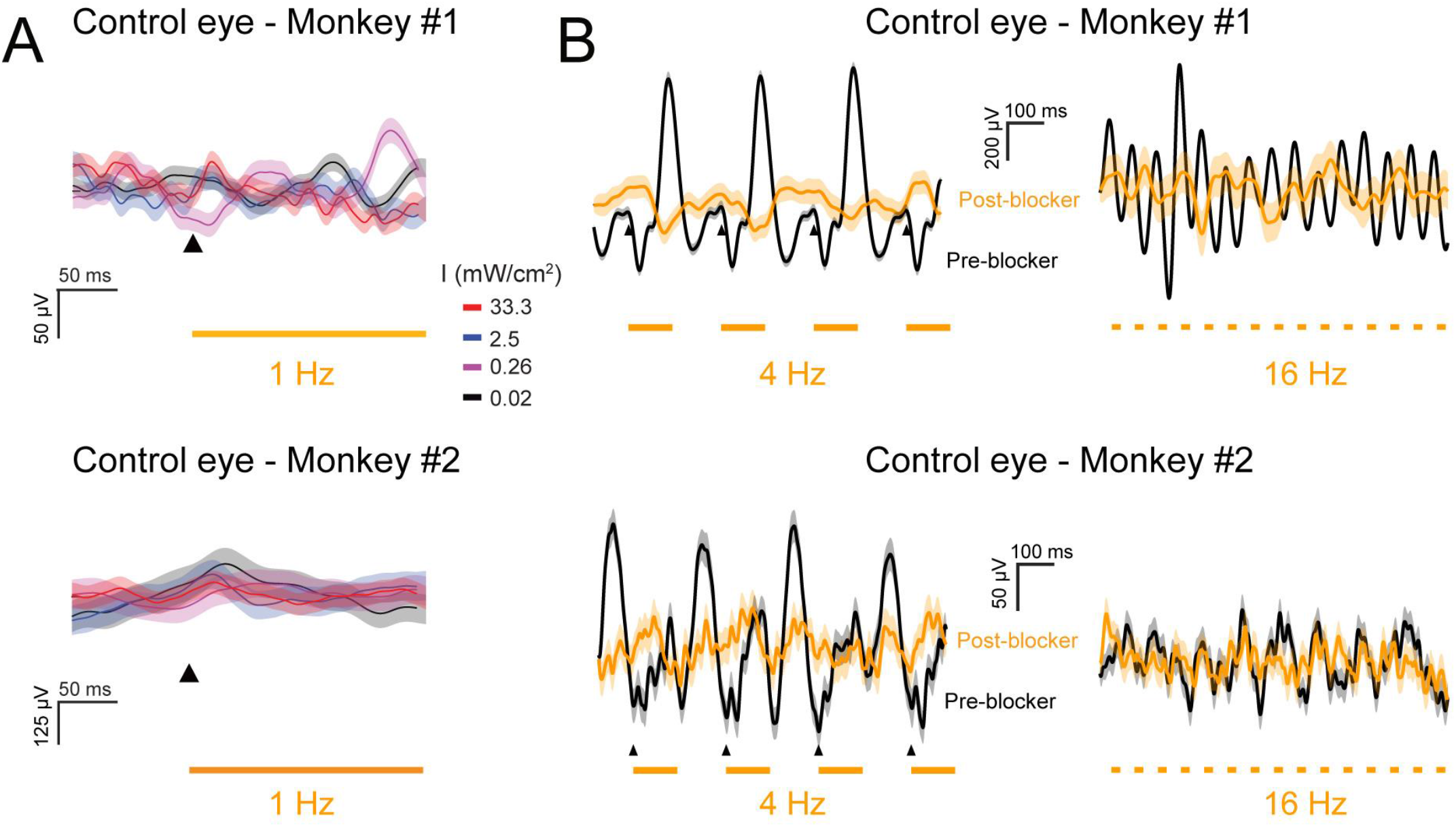
VEP recordings in the control eye. (A) VEP responses to the orange LED when the control eye was stimulated with light at different intensities (top: monkey M1 and bottom: monkey M2). We tested 4 different light intensities at a frequency of 1 Hz (33.3, 2.5, 0.26 and 0.02 mW/cm^2^), as indicated by the color code. (B) VEP responses to the orange LED when the control eye was stimulated at two different frequencies (4 and 16 Hz). Orange traces represent VEP traces recorded after the injection of synaptic blocker; black traces were recorded before the injection of synaptic blocker.

